# Predictive modeling of TMS-evoked responses: Unraveling instantaneous excitability states

**DOI:** 10.1101/2025.06.18.660274

**Authors:** Oskari Ahola, Lisa Haxel, Maria Ermolova, Dania Humaidan, Tuomas P. Mutanen, Mikael Laine, Matilda Makkonen, Elena Ukharova, Timo Roine, Pantelis Lioumis, Roberto Guidotti, Risto J. Ilmoniemi, Ulf Ziemann

## Abstract

Transcranial magnetic stimulation (TMS) combined with electroencephalography (EEG) and electromyography (EMG) provides a unique window into instantaneous cortical and corticospinal excitability states. We investigated 50 healthy participants to determine how fluctuations in pre-stimulus brain activity influence single-trial TMS-evoked potentials (TEPs) and motor-evoked potentials (MEPs). We developed a novel automated source-level TEP extraction method using individualized spatiotemporal priors that is robust against poor single-trial signal-to-noise ratios (SNRs) and ongoing oscillations. TEP and MEP amplitudes were predicted with linear mixed-effects models based on pre-stimulation EEG band-powers (theta to gamma), while accounting for temporal drifts (within-session trends), coil control, and inter-subject differences. We found that higher pre-stimulus sensorimotor alpha, beta, and gamma power were each associated with larger TEPs, indicating a more excitable cortical state. Increases in alpha and gamma power immediately before stimulation specifically predicted larger MEPs, reflecting increased corticospinal excitability. These results reveal relationships between ongoing oscillatory brain states and TMS response amplitudes, identifying EEG biomarkers of high- and low-excitability states. In conclusion, our study demonstrates the feasibility of single-trial source-level TMS–EEG analysis and shows that spontaneous alpha-, beta-, and gamma-band oscillations modulate motor cortical and corticospinal responsiveness. These findings pave the way for EEG-informed, brain-state-dependent TMS protocols to optimize neuromodulatory interventions in clinical and research applications.

## 1 INTRODUCTION

Combining transcranial magnetic stimulation (TMS) with electroencephalography (EEG) and electromyography (EMG) provides instantaneous information about cortical and corticospinal excitability states via TMS-related responses, such as TMS-evoked potentials (TEPs) and motor-evoked potentials (MEPs) (Ilmoniemi et al., 1997; Ilmoniemi and Kičić, 2010; Hallett, 2007). These neurophysiological measures reflect the responsiveness and plasticity of targeted cortical regions (Pellicciari et al., 2018; Gosseries et al., 2015; Voineskos et al., 2019) and are predictive markers of outcome in neurological and psychiatric disorders, including stroke and depression (Lefaucheur et al., 2020; Vucic et al., 2023; Strafella et al., 2023; Bembenek et al., 2020; Hordacre et al., 2019). Indeed, patients with a variety of brain network disorders who exhibit larger TMS-evoked responses often respond better to therapeutic interventions (Pellicciari et al., 2018; Bembenek et al., 2020; Strafella et al., 2023).

Despite their clinical relevance, TMS-evoked responses show substantial variability within and across individuals (Lioumis et al., 2009). This variability is partly driven by changing neuronal activity states (Voineskos et al., 2019) – as well as differences in structural connectivity (Casarotto et al., 2010). However, the specific electrophysiological features that determine whether the brain is in a high-or low-excitability state remain poorly understood. In particular, it is unclear how inhibitory and excitatory network dynamics before TMS influence the resulting TEP and MEP amplitudes. As a result, current standard TMS protocols ignore the momentary brain state and may deliver stimuli at suboptimal times, potentially compromising therapeutic success.

Recent evidence has begun to investigate how pre-stimulus EEG rhythms modulate TMS outcomes. Alpha-frequency oscillations were traditionally thought to index cortical inhibition, with higher alpha power suppressing excitability (Klimesch et al., 2007). Contrary to this notion, real-time EEG-triggered TMS studies have demonstrated pulsed facilitation of corticospinal excitability at specific phases of the sensorimotor *µ*-alpha-cycle, even finding that high alpha power can coincide with larger rather than smaller MEPs (Bergmann et al., 2019; Zrenner et al., 2018). Similarly, beta-frequency oscillations in motor cortex have been linked to the maintenance of the current sensorimotor state and GABAergic inhibitory tone, yet their exact relationship to excitability is complex (Groth et al., 2021). Gamma-band activity, reflecting local cortical excitatory–inhibitory circuit engagement, may conversely signal periods of heightened neuronal firing and readiness (Herrmann et al., 2010). Overall, these studies suggest that instantaneous oscillatory activity in multiple frequency bands could serve as a proxy for the brain’s excitability state, but a comprehensive, multi-band characterization of this relationship at the single-trial level is lacking. Furthermore, prior TMS–EEG studies often focused on sensor-level EEG or averaged responses, leaving open the question of how to reliably quantify single-trial TEPs from cortical sources.

To address these gaps, we present a methodology for single-trial source-level TEP analysis that accounts for the low spatiotemporal signal-to-noise ratio (SNR) of individual TMS–EEG trials. In the present study, we leverage this approach to explore the relationship between pre-stimulation EEG dynamics and subsequent TMS-evoked responses in a sample of 50 healthy adults. Specifically, we examine whether oscillatory power in canonical frequency bands prior to stimulation can predict the amplitude of the induced TEPs in the targeted cortex and the corresponding MEPs in the periphery. By identifying robust EEG markers of instantaneous cortical and corticospinal excitability, our work aims to deepen the neurophysiological understanding of TMS response variability. Importantly, these insights also lay groundwork for brain-state-dependent TMS protocols, whereby stimulation can be timed to an optimal excitability state to enhance neuromodulatory effects. Such EEG-informed TMS strategies hold promise for increasing the efficacy of therapeutic and rehabilitative interventions across a range of brain disorders.

## 2 METHODS

We conducted a study investigating TMS-evoked EEG and motor responses at two European research centers in the ConnectToBrain synergy project funded by the European Research Council (grant no. 810377). The experimental protocol was standardized while accommodating for site-specific equipment variation.

### 2.1 Data acquisition

The measurements were performed by two research groups: the Brain Networks & Plasticity Laboratory at the Hertie Institute for Clinical Brain Research, University of Tübingen, Germany, and the TMS Group at the Department of Neuroscience and Biomedical Engineering, Aalto University, Finland.

The combined dataset consists of 50 healthy right-handed participants (28 women) of ages 28 ± 6 years (mean ± standard deviation). The TMS–EEG measurement contained four blocks of 300 biphasic single pulses at an intensity of 110% of the resting motor threshold (rMT) targeted at the left motor hotspot with a randomized inter-trial interval of 4.0–4.5 or 2.5–3.5 s, in Aalto and Tübingen, respectively. RMT was defined as the minimum stimulation intensity that resulted in MEPs with a minimum peak-to-peak difference of 50 *µ*V in at least 50% of trials (Rossini et al., 2015).

The TMS–EEG data were recorded with TMS-compatible NeurOne 128-and 64-channel EEG systems (Bittium Ltd, Finland) with a sampling rate of 5 kHz. EMG was recorded from the right abductor pollicis brevis (APB) and the first dorsal interosseus (FDI) muscles in a bipolar belly–tendon montage. TMS was administered with a Nexstim NBS 5.2.4 or NBT 2.2.4 (Nexstim Plc, Finland) system with a Nexstim cooled coil (Aalto) or with a MagVenture (MagVenture Inc, United States) R30 or X100 stimulator with a Cool-B65 coil (Tübingen). Neuronavigation was performed using individual T1-weighted magnetic resonance images (MRIs) with Nexstim or Localite (Localite GmbH, Germany) systems in Aalto and Tübingen, respectively. The TMS click sound was masked with individually calibrated noise (Russo et al., 2022) using ER-3C insert earphones (Etymotic Research Ltd, United States).

T1- and T2-weighted MRIs were acquired with magnetization-prepared rapid acquisition gradient echo (MPRAGE) and turbo spin echo (TSE) sequences, respectively, using 3T Siemens Skyra and Prisma scanners (Siemens Plc, Germany). For further specification on experimental details, see Section 1 in the Supplementary Materials.

All subjects gave their written informed consent for participating in the study, which was approved by the ethics committees of the University of Tübingen (810/2021BO2) and the Helsinki University Hospital (HUS/1198/2016). The study was conducted in compliance with the Declaration of Helsinki.

### 2.2 Preprocessing

The TMS–EEG data were preprocessed with Matlab 2024a (The MathWorks Inc, United States) using custom scripts and EEGLAB 2024.2 (Delorme and Makeig, 2004) with the TESA toolbox (Rogasch et al., 2017; Mutanen et al., 2020) in an automated fashion. The pre- and post-stimulus periods, which were initially defined to be [−1250, −25) and [−20, 300) ms relative to the TMS pulse, respectively, were processed separately to not violate causality (*i*.*e*., induce data leakage). Note that our post-stimulus-focused data includes times from the official pre-stimulus period (*i*.*e*., times before TMS), but these times are not included in the pre-stimulus-focused period that is later used to predict post-stimulus activity.

We interpolated TMS pulse artifacts and removed noisy channels by separately considering both periods. We then calibrated independent component analysis (ICA) (Hyvärinen and Oja, 2000; Pion-Tonachini et al., 2019) filters from pre-stimulus, 2-Hz highpass filtered, and average referenced data to suppress ocular artifacts. The calibrated filters were subsequently applied to non-filtered average-referenced pre- and post-stimulus periods. This way of applying ICA also mitigates suppressing TEP-related neuronal activity that could be mixed in the ICA filters if the ICA decomposition was derived from both pre- and post-stimulus periods.

For the pre-stimulus period, we reconstructed bad channels via spherical spline interpolation, applied 4^th^-order zero-phase Butterworth bandpass and powerline bandstop filtering (2–90, and 48–52 Hz, respectively), downsampled the data to 1000 Hz, and cropped the data to [−1045, −45) ms. We then rejected trials based on standardized standard deviation statistics of the pre-stimulus period.

For the post-stimulus-focused period, we suppressed noise and reconstructed contaminated channels using the source-estimate-utilizing noise-discarding (SOUND) algorithm (Mutanen et al., 2018). We then suppressed muscle artifacts with the signal-space-projection-source-informed reconstruction (SSP–SIR) algorithm (Mutanen et al., 2016), and re-interpolated an extended TMS pulse artifact window. After cropping the post-stimulus period to [−20, −150) ms, we removed bad trials based on standardized standard deviation statistics of the post-stimulus period. The post-stimulus baseline correction range was [−20, −10] ms. The pre-and post-stimulus data were analyzed in average reference. Contrary to common practices, we do not downsample or directly bandpass filter our post-stimulus data to avoid additional temporal smearing and edge effects.

EMG preprocessing included baseline correction, linear detrending, and pre-innervation-based trial exclusion within the pre-stimulation window of [−100, −10] ms. We extracted MEP amplitudes as the peak-to-peak (*i*.*e*., minimum-to-maximum voltage difference) from [20, 50] ms of each trial from the channel with the average higher amplitude. Detailed preprocessing specifications on channel and trial rejection, hyperparameters, baseline corrections, and intermediate referencing and time range selections, are provided in Section 2 in the Supplementary Materials.

### 2.3 TEP extraction

Equivalent current dipoles can be used to represent small source-current concentrations (Henderson et al., 1975; Sarvas, 1987). The TMS-evoked potentials (TEPs) for each trial were modeled by the amplitude of time-constrained, fixed-position, fixed-orientation current dipole fits. When fitting dipoles to single-trial data, we used prior temporal, spatial, and orientational information from the average response to mitigate the effect of poor singletrial SNR and ongoing oscillations. The best-fitting dipoles in the average response were automatically identified using pseudoinverse by maximizing the coefficient of determination (*R*^2^) and response stability.

For each response, we initially defined the following time ranges **T**: N15: [12, 25], P30: [25, 40], N45: [40, 55], and P60: [55, 73] ms. Fitting was done in reverse temporal order. To avoid response overlapping, the upper edge of each time window prior to P60 was sequentially adjusted to ensure a minimum 5-ms gap from the minimum time of the subsequent response.

For each response, we defined candidate dipoles 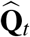 at positions 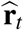 for each time point *t ∈* **T** as

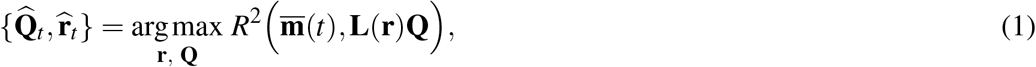

where **Q** is the dipole moment, 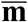 is trial-averaged EEG data **m**, *t* is the time, and **L**(**r**) is the lead field at position **r**.

For each dipole 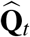 at 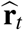, we defined activation time ranges 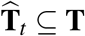 around *t* as continuous intervals during which the dipole fixed at 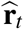 had at least 90% of the *R*^2^ and deviated no more than 10 degrees relative to 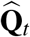. with an unit-length orientation 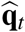.To prioritize long intervals while mitigating the effect of prolonged low-amplitude dipolar (resulting in sufficient *R*) activity with little spatial variability, the optimal fitting time range 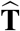 for the response was then defined as the interval,

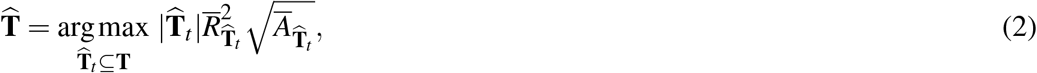

where 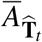 (in units of nAm) and 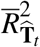 are the average dipole amplitude and *R*^2^, respectively, in time range 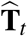. The optimal dipole position 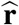 and orientation 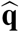 correspond to this time window.

Finally, for each trial *i*, the dipole fixed at 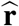 with orientation 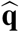 in time range 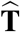 was selected as

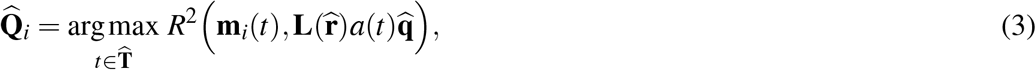

The scalar amplitude *a*(*t*) — obtained by solving 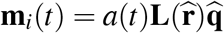 using least squares — of a best-fitting fixed-orientation dipole 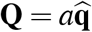, can be negative due to polarity reversal. We use the absolute amplitude to represent the magnitude of cortical excitation. To mitigate artifactual effects and to address poor single-trial SNR, for each response, we only included subjects whose optimal dipole fit to the average response (resulting in time range 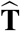) had an *R* of at least 75% and an amplitude of at least 20 nAm. The extraction procedure is illustrated in Fig. 1.

**Figure 1.**
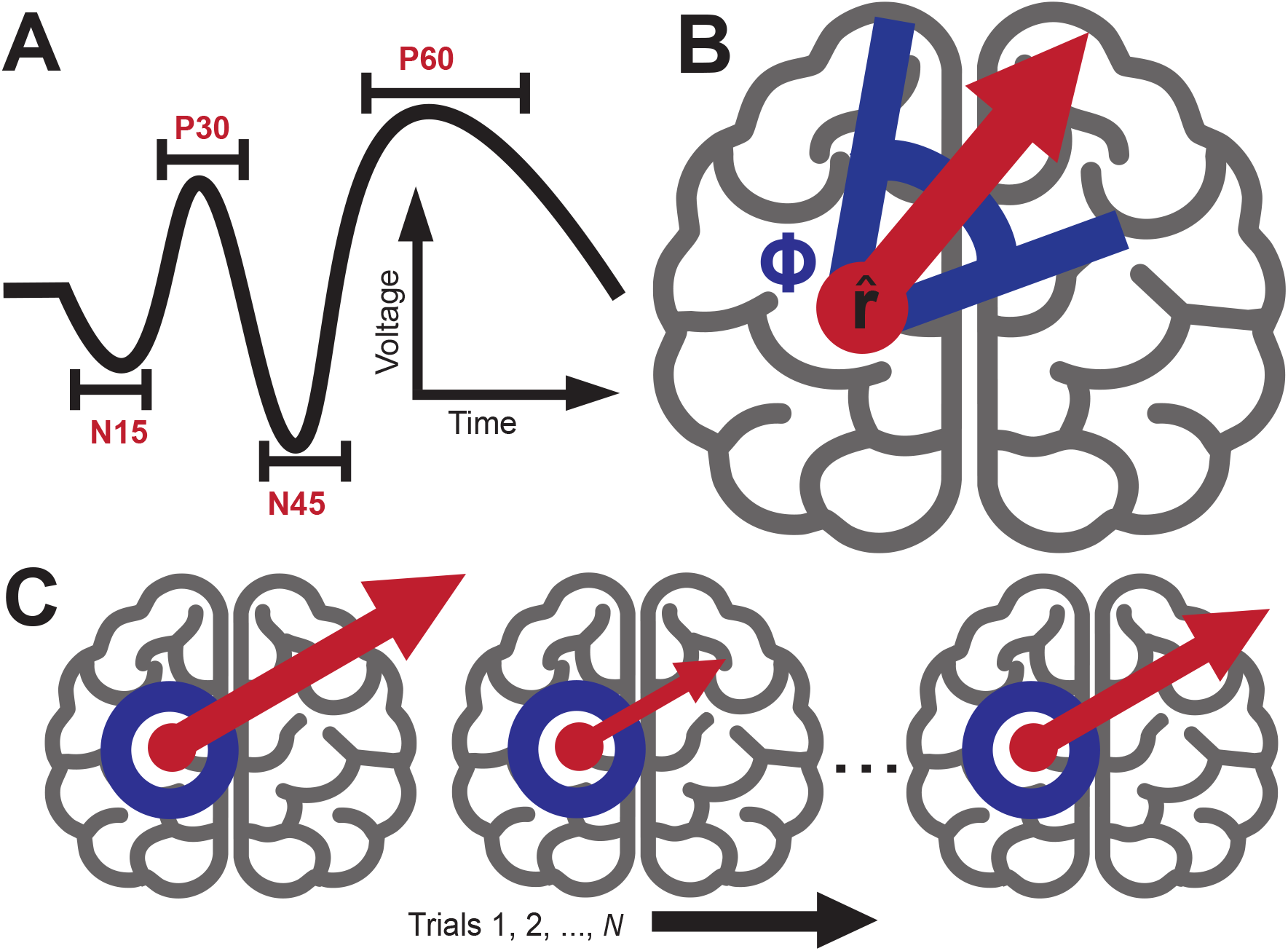
TEP extraction procedure. **A**: Illustration of TEP components with fitting time ranges determined by maximizing dipole coefficient of determination (*R*^2^), amplitude, and orientation stability. The fitting range was adjusted to ensure a minimum 5-ms gap to the subsequent response. **B**: Determination of an extraction time range 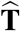 for a response where dipole orientations fitted to the dipole position 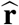 deviated less than *ϕ*= 10 degrees and had an *R*^2^ of at least 90% relative to the optimal dipole. **C**: Fixed-orientation dipoles with different amplitudes modeling single-trial responses fitted to fixed positions 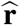. The figure was created with Adobe Illustrator using the Text to Vector graphics tool for the brains.

### 2.4 Feature extraction

We performed source-space analysis using subject-specific MRIs for the pre-stimulation period [−1045, −45) ms using MNE Python 1.7.0 (Gramfort et al., 2013) and custom scripts. Source estimates were acquired with minimum-norm estimation (MNE) (Hämäläinen and Ilmoniemi, 1994), projected onto the *fsaverage* brain, and parcellated according to the Desikan– Killiany atlas (Desikan et al., 2006). To investigate effects near response and stimulation locations, source-space signals were also extracted from custom parcels defined as source-point clusters with maximal geodesic distances of 2 cm to the nearest source positions of a) the individual response (dipole) locations and b) the MNI-Talairach left motor hand knob at (−40, −20, 62) mm.

For each parcellated pre-stimulus time series, we calculated power spectral densities (PSDs) for the theta, alpha, beta, and gamma frequency bands with the multi-taper method. The frequency bands were determined individually based on individual peak alpha-power frequency (Klimesch, 1999), identified from the average pre-stimulus sensor space PSD. This resulted in average frequency ranges of theta = 3.8–7.3, alpha = 7.3–12.3, beta = 12.3–32.3, and gamma = 32.3–82.3 Hz.

For each trial, the coil control (stimulation accuracy) value was computed as a weighted average of the differences between the actual and target stimulation parameters — specifically position (in units of mm), normal, and direction (in degrees). To emphasize deviations in coil position (which tend to be more variable than orientation) and to reduce possible multicollinearity, we applied weights of 0.7 for position and 0.15 for normal and direction. Missing coil control data (6.2% of trials) was handled with median replacement. Further information on feature extraction is available in Section 4 in the Supplementary Materials.

### 2.5 Predictive modeling

We predicted log-transformed absolute dipole (source-localized TEP) and MEP amplitudes using linear mixed-effect models (LMMs) (Lindstrom and Bates, 1988; Seabold and Perktold, 2010) with subject-level random effects as

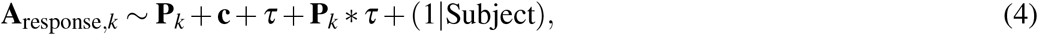

where **P**_*k*_ are the band-powers of each frequency band (theta, alpha, beta, and gamma) in parcel (brain region) *k* and **c** is the coil control for each trial (weighted pose deviances from the target). *τ* is the maximum-normalized post-processed trial number for each subject (proxy for time, referred as such from here onwards), and **P**_*k*_ *\* τ* denotes the interaction between band-power and time.

We log-transformed power, time, and coil control parameters and standardized them zero-mean and unit variance. Powers were within-subject standardized to account for subject-specific baselines and variability, and the time and coil control were globally standardized. While our primary focus is the effect of fluctuations in power, we included the coil control as a covariate to account for targeting accuracy — a known confound in TMS. Additionally, we included band-power–time interactions to distinguish instantaneous fluctuations from over-time trends that may exist between power-based predictors and response amplitudes.

### 2.6 Statistics

The two-tailed *p*-values of the LMM coefficients were determined using Wald *t*-tests. The *p*-values were corrected for multiple comparisons with the Bonferroni method while excluding the model intercept (Seabold and Perktold, 2010). All *p*-values of LMM coefficients reported in this text have been Bonferroni-corrected. LMM goodness-of-fits were evaluated using conditional and marginal *R*^2^ (Nakagawa and Schielzeth, 2013); see Section 5.2 in the Supplementary Materials.

The two-tailed *p*-values of Spearman correlation coefficients were computed using the t-distribution under the null hypothesis of no monotonic relationship (Virtanen et al., 2020).

## 3 RESULTS

### 3.1 Predictive features

We found several predictors of TEP and MEP amplitude across multiple parcels (brain regions). The strongest positive TEP predictors were at the alpha, beta, and gamma bands and spatially similar across especially temporally adjacent responses. TEPs with longer latencies (N45 and P60) were more positively affected by changes in bilateral centro-parietal beta and bilateral sensorimotor gamma power, whereas the earlier TEPs (N15 and P30) were more affected by bilateral sensorimotor or parietal alpha power. P30 was also positively modulated by frontal beta power.

MEPs were most positively affected by ipsilateral (on the side of stimulation) sensorimotor alpha-frequency (similar to the sensorimotor *µ*-rhythm) and bilateral frontal gamma power. These findings align with previous research (Thies et al., 2018; Wischnewski et al., 2022; Haxel et al., 2025).

We compared absolute predictive coefficients of the models from the custom and anatomical parcels (see Section 2.4) to compare local effects at regions affected by stimulation. At the custom motor hand-knob parcel, the predictive alpha power coefficient for MEPs was the at the 100^th^ percentile (*β* = 0.07) and at the 97^th^ percentile for N15 (*β* = 0.04), and the gamma power coefficient of P60 was at the 97^th^ percentile (*β* = 0.04). Gamma power yielded similar effects at the response-location derived parcels for N15, P30, and N45 (*β* = 0.03; 100^th^, 97^th^, and 98^th^ percentiles, respectively), and P60 (*β* = 0.04; 100^th^ percentile). Similarly, the beta power coefficient at the response-location derived parcel was at the 97^th^ percentile for N45 (*β* = 0.02) and at the 98^th^ percentile for P60 (*β* = 0.03).

The band-power coefficients for each response on each anatomical label are displayed in Fig. 2.

**Figure 2.**
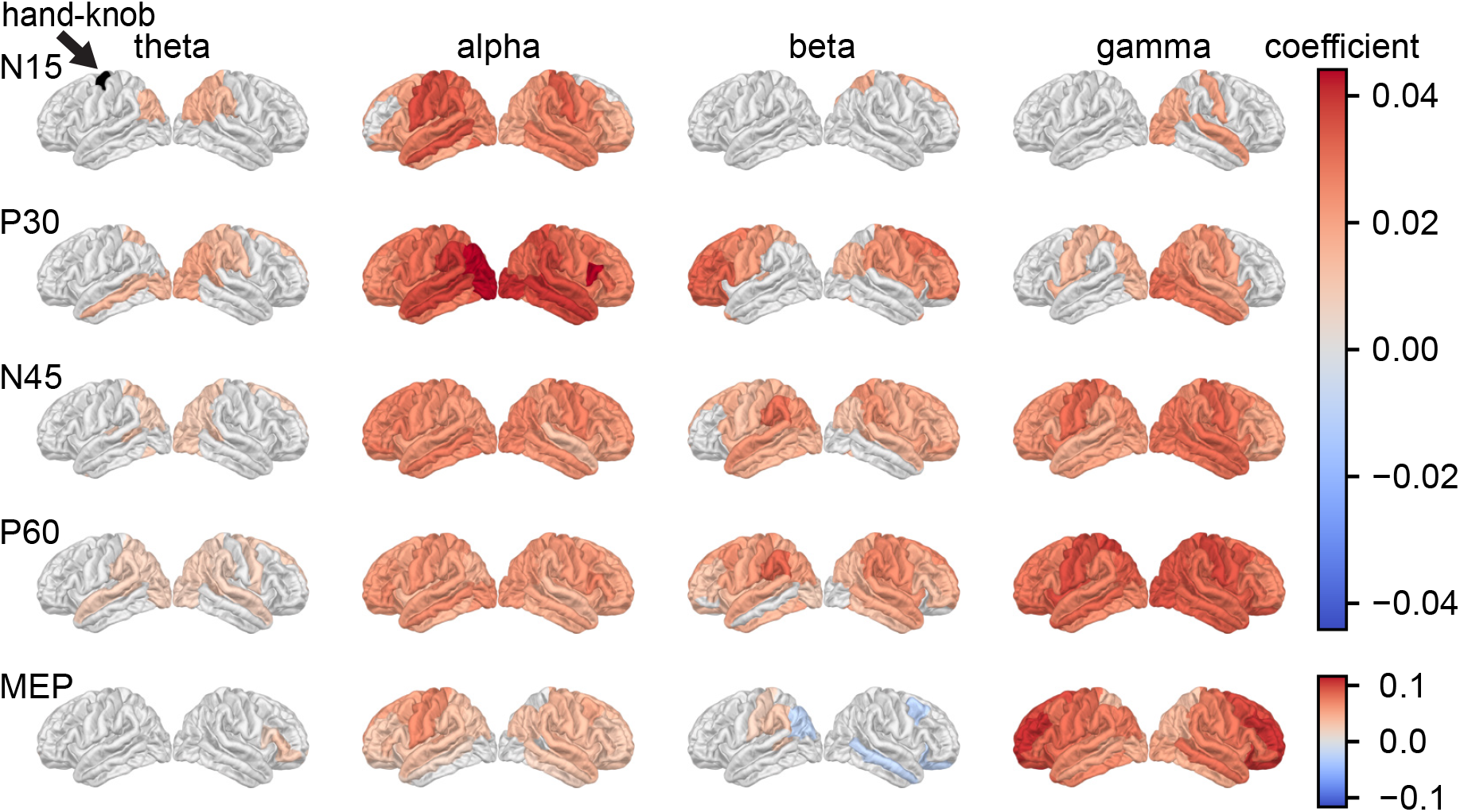
Band-power coefficients of linear mixed-effects models. Each anatomical parcel represents the frequency band-power coefficient of the model of the respective parcel modeling the respective response. Gray regions indicate insignificant (*p ≥* 0.05; Bonferroni corrected) coefficients. The predictors were scaled to zero mean and unit variance. The frequency ranges were derived based on individual alpha peaks (9.7 ± 1.2 Hz), resulting in average ranges of theta = 3.8–7.3, alpha = 7.3–12.3, beta = 12.3–32.3, and gamma = 32.3–82.3 Hz (see Section 4 in the Supplementary Material for more information). Note that the colorbar scale is different for TEPs and MEPs. The motor hand-knob parcel is indicated as the black region in the left uppermost brain. Across anatomical parcels, maximum model conditional and marginal *R*^2^ (in %), were, respectively, N15: 70.5 and 0.5, P30: 50.3 and 0.6, N45: 51.2 and 0.6, P60: 34.7 and 1.2, and MEP: 42.8 and 2.1. Note that the number of subjects differs for each response type, which makes the results less comparable. Supposing that other variables are held constant, a coefficient of *β* implies that a 1 SD change in the log-transformed predictor has an exp(*β*) effect on the non-logged outcome variable, *e*.*g*., exp(0.1) − 1 *≈* 10.5%.

The ongoing measurement time (*i*.*e*., trends; modeled with trial numbers) was a positive predictor for MEPs, with a maximum coefficient strength of *β* = 0.11 (*p<* 0.05). The effect was not notable for TEPs with a maximum of *β* = 0.02 (*p<* 0.05) for N15. The band-power– time interaction terms yielded smaller coefficients: |*β*| *<* 0.03 for TEPs and |*β*| *<* 0.04 for MEPs (both *p<* 0.05). Most notably, MEP was negatively modulated by gamma power trends and positively by alpha power trends. Visualization is provided in Section 5.1 of the Supplementary Materials.

The coil deviations (subject means; position: 2.2 ± 1.0 mm, direction: 1.3 ± 0.7, and normal: 1.3 ± 0.8 degrees) had small effects on N15 (minimum of *β* = −0.04) and larger maximal negative effect on MEPs (*β* = −0.14); both *p<* 0.05.

### 3.2 Response stability and correlations

Stability across different outcome measures provide essential information on inter-response relationships, and consequently, differences between cortical and corticospinal excitability. Assessing these factors increases overall fidelity of the modeled low SNR single-trial TMS– EEG responses.

We observed particularly high across-trial variability in dipole (source-localized TEP) amplitudes and goodness-of-fits (*R*^2^ values). The key characteristics across trials are displayed in Table 1 and Fig. 3. Analysis of the relationship across TEPs and MEPs revealed significant positive correlations across all responses except for N15 (Fig. 3). Notably, TEP–TEP correlations were stronger between temporally adjacent responses compared to TEP–MEP correlations. The amplitudes across trials were always positively correlated with the goodness-of-fit (Fig. 3) with an across-subject standard deviation of 0.14 for N15 and under 0.1 for the subsequent TEPs.

**Table 1.** Response characteristics across MEPs and TEP components fitted to single trials. Values are presented as mean ± standard deviation (STD). Parameters with “mean” and “STD” are within-subject means and STDs. *N*: Number of included subjects whose dipole fitted to the average response had an *R*^2^ of at least 75% and an amplitude of at least 20 nAm (not the average dipole fit across trials, which is displayed in the Table). Amplitude: Absolute value of dipole strength (non-scalar amplitude). Dipole PTP (peak-to-peak): Sensor-space PTP of the forward-projected dipole. PTP data: Sensor-space PTP at the time of the dipole (or the MEP PTP) and ΔPTP is the PTP difference between the forward predicted dipole and sensor-space PTP. The PTPs are calculated as min-max differences. Latency: Post-stimulus latency of the optimal dipole fit. Window size: Width of the fitting time range. See Table 3 in the Supplementary Materials for average responses.

| Parameter | N15 | P30 | N45 | P60 | MEP |
| --- | --- | --- | --- | --- | --- |
| **Fit Overview, *N*=** | 11 | 23 | 41 | 45 | 50 |
| $R^2$ mean (%) | 61.5 ± 14.5 | 44.9 ± 14.9 | 49.6 ± 12.8 | 55.5 ± 13.9 | - |
| $R^2$ STD (%) | 19.1 ± 4.4 | 22.1 ± 2.8 | 22.9 ± 2.3 | 21.9 ± 3.1 | - |
| Amplitude mean (nAm) | 124.3 ± 140.5 | 87.0 ± 73.5 | 129.4 ± 95.3 | 106.0 ± 46.2 | - |
| Amplitude STD (nAm) | 58.0 ± 76.3 | 47.3 ± 36.5 | 63.8 ± 44.7 | 49.3 ± 25.9 | - |
| Dipole PTP mean (μV) | 16.0 ± 6.8 | 13.9 ± 8.6 | 18.9 ± 7.7 | 19.0 ± 5.8 | - |
| Dipole PTP STD (μV) | 7.0 ± 2.4 | 7.4 ± 3.8 | 9.3 ± 3.8 | 8.7 ± 2.6 | - |
| PTP data mean (μV) | 21.6 ± 8.4 | 22.5 ± 10.3 | 29.5 ± 9.7 | 28.8 ± 7.8 | 1144 ± 1030 |
| PTP data STD (μV) | 7.4 ± 2.8 | 8.5 ± 3.4 | 10.4 ± 3.6 | 10.1 ± 3.1 | 906 ± 641 |
| ΔPTP mean (μV) | 5.7 ± 3.5 | 8.7 ± 4.2 | 10.7 ± 4.2 | 9.9 ± 4.0 | - |
| ΔPTP STD (μV) | 4.2 ± 2.2 | 6.2 ± 2.8 | 7.5 ± 2.8 | 6.7 ± 2.5 | - |
| Latency mean (ms) | 16.7 ± 1.9 | 32.8 ± 3.5 | 47.4 ± 2.2 | 62.8 ± 3.5 | - |
| Latency STD (ms) | 2.4 ± 0.5 | 1.7 ± 0.6 | 2.5 ± 0.7 | 4.0 ± 1.2 | - |
| Window size (ms) | 7.7 ± 1.4 | 5.1 ± 1.7 | 7.5 ± 2.3 | 12.2 ± 4.0 | 30.0 |

**Figure 3.**
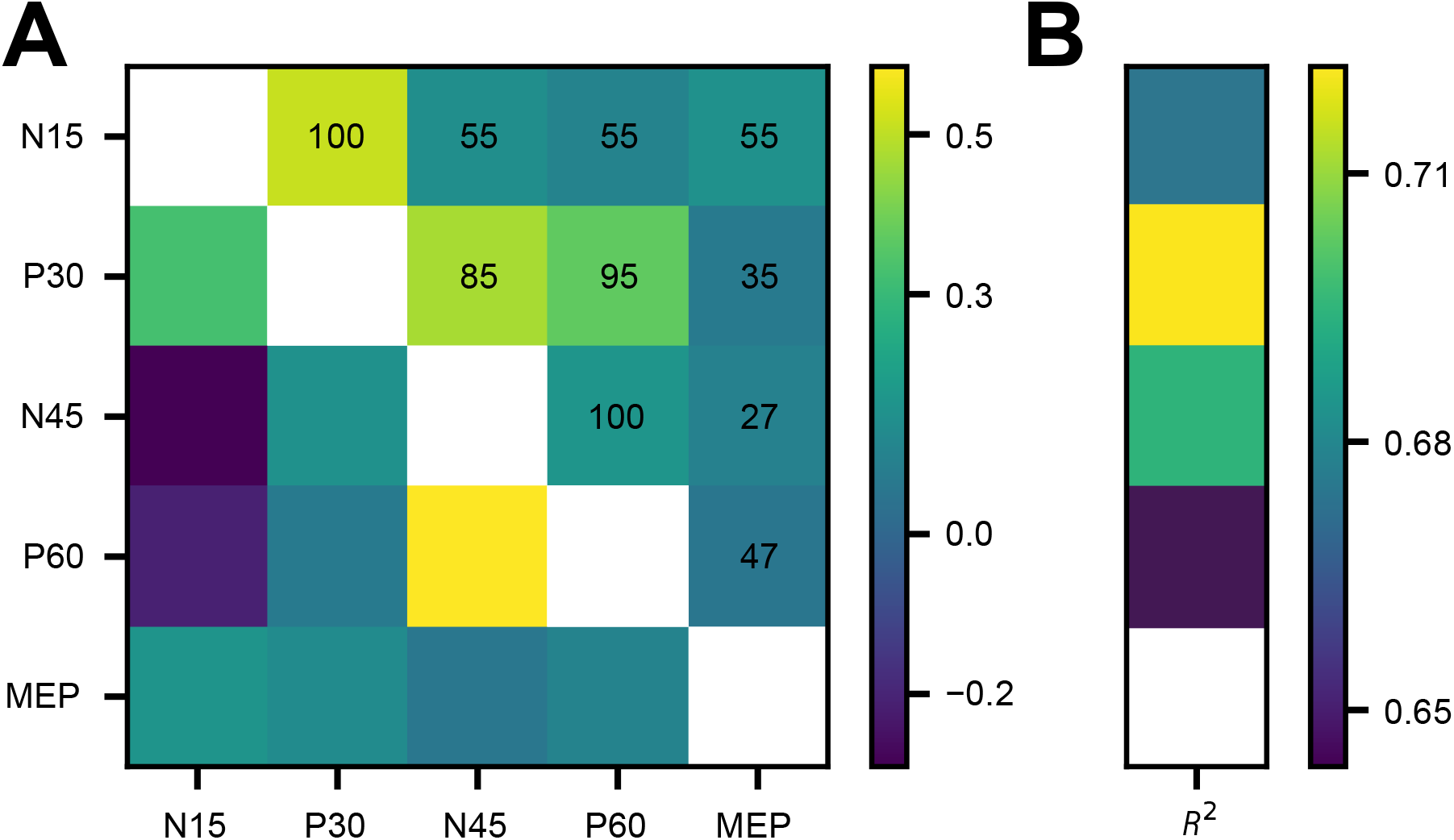
Spearman correlation analysis of response amplitudes. **A**: Lower triangle: Mean, and upper triangle: Standard deviation of significant (*p<*0.05) correlation coefficients of response amplitudes across trials. TEP–TEP correlations are calculated with scalar amplitudes and TEP–MEP correlations with absolute amplitudes. The percentage of subjects with significant correlations is displayed in the respective cell. **B:** Mean correlations between absolute TEP (dipole) amplitudes and dipole *R*^2^ (always significant; *p<*0.05). Only subjects reported in Table 1 are included in the analysis.

## 4 DISCUSSION

### 4.1 Principal advances

The present work introduces a single-trial, source-level extraction method for TMS-evoked potentials and demonstrates that moment-to-moment fluctuations in pre-stimulus EEG band-powers systematically predict both cortical (TEP) and corticospinal (MEP) response amplitudes. By extracting single-trial TEPs from their dominant dipolar sources and modeling amplitudes with linear mixed-effects, we show that high-excitability states of the motor cortex are signaled by elevated sensorimotor alpha, beta, and gamma power, whereas low-excitability states appear when these rhythms are weak. These findings establish a mechanistic link between ongoing oscillations and TMS responsiveness and provide a quantitative basis for real-time, brain-state-dependent neuromodulation (Bergmann, 2018; Humaidan et al., 2024).

### 4.2 Oscillatory markers of excitability

#### Sensorimotor *µ*-alpha rhythm

We observed that greater alpha power in the left sensorimotor cortex and other brain regions predicted larger TEPs and MEPs. Although alpha-frequency activity is often interpreted as an inhibitory “idling” rhythm, a recent real-time EEG-triggered TMS study has shown facilitation of corticospinal output during high-power or alpha states (Bergmann et al., 2019). One explanation is that synchronous alpha activity represents a low-noise “readiness” state in which neurons require less additional input to reach firing threshold. This view also fits with cognitive work linking alpha to efficient information gating during memory encoding and retrieval (Klimesch, 1999).

#### Beta and gamma rhythms

Increases in beta and gamma power were positive predictors for later-latency TEP amplitudes, whereas prefrontal gamma power uniquely predicted MEP size. These rhythms are closely coupled to local excitation–inhibition balance; fast beta and gamma oscillations in the motor cortex scale with intracortical GABAergic inhibition and learning capacity (Rossiter et al., 2014; Rempe et al., 2022; Zich et al., 2025). Elevated beta and gamma activity may also mark a predictive or preparatory network state in which pyramidal neurons are synchronously primed for external perturbation — consistent with the idea that participants implicitly anticipate both the TMS click and the ensuing finger twitch.

#### Network-level influences

Beyond the stimulated primary motor cortex, we identified frontal gamma sources that modulated MEP amplitudes, suggesting that distributed sensorimotor and premotor circuits shape local excitability. This aligns with evidence that large-scale oscillatory networks predict corticospinal output (Ermolova et al., 2024; Haxel et al., 2025). Together, the data support a model in which instantaneous excitability is determined not only by local oscillations but also by the broader functional network state.

### 4.3 Technical confounds: coil control and hotspot fidelity

Coil deviations exerted a markedly greater influence on MEPs than on TEPs, because even small spatial errors shift the induced current away from the optimal corticospinal hotspot, whereas a distributed cortical response can still be evoked at nearby sites. Furthermore, is likely that the effect of coil deviations is larger in reality as the mapped hotspot and the deviations from the mapped hotspot can result in higher or lower effective stimulation of the true hotspot, resulting in higher or lower MEP (or TEP) amplitudes, respectively. The effect of coil control and hotspot fidelity could both be addressed by robotic control and machine learning-based MEP or TEP hotspot mapping (Matsuda et al., 2024; Nieminen et al., 2022; Tervo et al., 2022).

### 4.4 Methodological strengths and limitations

Our single-trial pipeline leveraged spatiotemporal priors from the grand-average TEP to constrain dipole location and orientation, thereby reducing the danger of overfitting to the low-SNR post-pulse interval and to ongoing oscillations. The approach yielded (*i*) significant scalar amplitude TEP–TEP correlations across contiguous trials, (*ii*) systematic relationships between pre-stimulus power and response amplitude, and (*iii*) spatially focal predictors that peaked at the stimulation and response sites — all hallmarks of genuine physiological signal rather than noise.

Alternative strategies — *e*.*g*., fitting free-orientation dipoles or expanding to multi-source models — can improve numerical goodness-of-fit but risk “chasing” noise or volume-conducted ongoing oscillations. We therefore prioritized specificity over raw fitting accuracy in exchange for cleaner neuroanatomical interpretability. However, while fixing the dipole orientation to the dominant orientation suppresses the contribution of other sources, it also limits the effects of true TEP orientation variability at the location of interest. A pragmatic future compromise could involve pooling early TEP features across two or three consecutive trials to boost SNR without sacrificing temporal resolution.

Residual TMS-related artifacts remain an inevitable concern. Despite the use of SOUND (Mutanen et al., 2018) and SSP–SIR (Mutanen et al., 2016), these artifacts may still contaminate early TEPs. Additionally, trials with high-frequency power pre and post TMS can also indicate the increased presence of artifacts and noise. This could lead to non-plausible dipoles with high-amplitudes, possibly distorting the effect between high-power modulations and response amplitude. We therefore advise combining subject-specific stimulation sites that elicit high-SNR TEPs (Casarotto et al., 2022; Mutanen et al., 2013) with adaptive, per-subject preprocessing parameters in future work.

### 4.5 Toward real-time, brain-state-dependent stimulation

Because the fixed-effects predictors explained only a modest share of variance (marginal *R*^2^ *≈* 1–2%), closed-loop algorithms that continuously maximize response amplitude — rather than relying on static thresholds — are likely to outperform simple EEG biomarkers. Our source-level TEP read-out offers a rich feedback channel reflecting cortical excitability and is not confounded by spinal factors that affect MEPs. A possible pipeline would be

1. **Calibration:** Collect 100 open-loop trials to fit individualized ICA, SOUND, and SSP–SIR filters, acquire TEP estimates, and oscillatory predictors.
2. **Online tracking:** Monitor alpha, beta, and gamma power at the stimulation site, at its frontal network nodes, and at the site of the TEP.
3. **Adaptive scheduling:** Deliver the next pulse at the predicted peak-excitability window — or postpone it if a low-excitability window is detected.
4. **Continuous re-learning:** Update the filter and model weights every 30 trials to account for slow drifts (time-on-task “trends”) and fatigue.

## 5 CONCLUSIONS

Our results confirm that single-trial, source-resolved TEPs can be reliably extracted and that fluctuations in alpha, beta, and gamma frequency power shortly before stimulation are reliable signatures of motor cortical and corticospinal excitability. In practical terms, these findings pave the way for EEG-informed, brain state-dependent TMS as a route to maximize neuromodulatory impact — both inside and beyond motor cortex.

## 6 CREDIT AUTHORSHIP CONTRIBUTION STATEMENT

**Oskari Ahola:** Writing – review and editing, Writing – original draft, Visualization, Software, Validation, Methodology, Investigation, Formal analysis, Data curation, Conceptualization. **Lisa Haxel:** Writing – review and editing, Visualization, Software, Conceptualization. **Maria Ermolova:** Writing – review and editing, Investigation, Data curation. **Dania Humaidan:** Writing – review and editing, Investigation. **Matilda Makkonen:** Writing – review and editing, Investigation, Data curation. **Mikael Laine:** Writing – review and editing, Investigation. **Tuomas P. Mutanen** Writing – review and editing, Investigation. **Elena Ukharova:** Writing – review and editing, Investigation, Data curation. **Timo Roine:** Writing – review and editing, Investigation. **Pantelis Lioumis:** Writing – review and editing, Investigation, Supervision. **Roberto Guidotti:** Writing – review and editing. **Risto J. Ilmoniemi:** Writing – review and editing, Project Administration, Funding acquisition. **Ulf Ziemann:** Writing – review and editing, Project Administration, Funding acquisition, Supervision, Conceptualization.

## 7 DATA AVAILABILITY

De-identified data is available from the corresponding authors upon reasonable request, subject to ethical approval and data agreements.

## 8 CODE AVAILABILITY

The main code used in this study is publicly available and can be accessed at https://github.com/OAhola/TEP_MEP_Predictions.

## 10 DECLARATION OF GENERATIVE AI AND AI-ASSISTED TECHNOLOGIES IN THE WRITING PROCESS

During the preparation of this work the author(s) used ChatGPT and Gemini in order to make refinements to the text. After using this tool, the author(s) reviewed and edited the content as needed and take(s) full responsibility for the content of the published article.

## 11 ACKNOWLEDGEMENTS

This study has received funding from the European Research Council (ERC) under the European Unizon’s Horizon 2020 research and innovation programme (Grant agreement No. 810377). We thank Maxim Schaier, Aishwarya Bagal, Athea Laske, Hailin Wang, Olivier Roy, David Vetter, Noora Kainulainen, Ida Granö, Maria Nazarova, Shokoofeh Parvin, Ana Soto, Ida Ilmoniemi, Timo Tommila, and Ilkka Rissanen for assistance in TMS–EEG measurements, Joonas Laurinoja for measurements assistance and protocol structuring in Aalto, and Miriam Kirchhoff and David Vetter for discussions on phase-effects. We also thank Pedro Gordon and Andreas Jooß for assistance in MRI data collection. We additionally thank Prof. Gian Luca Romani, Prof. Laura Marzetti, Prof. Vittorio Pizzella, Delia Lucarelli, and Giulia Pieramico for feedback on the text.

