## Supplementary material for "Predictive modeling of TMS-evoked responses: Unraveling instantaneous excitability states"

### 1 MEASUREMENT PARAMETERS

**Table 1. Site-specific measurement parameters.** A: The first 12 measurements were conducted at the BioMag Laboratory and the latter 11 at Aalto University using different Nexstim systems (in the main text, we refer to these experiments as ones performed at Aalto). B: R30 was used for the first 15 participants and X100 for the latter 12. The stimulator intensities of R30 and X100 are relatively comparable because they are calibrated to the same output power. The same holds for the used Nexstim systems. C: Uniform distribution in Aalto and triangular in Tübingen. MSO: Maximal stimulator output, ITI: Inter-trial interval, and LP: low-pass. The rMT was defined as the stimulation intensity exceeding 50 $\mu$V in at least 50% of at least 10 trials. Current and filter refer to the NeurOne system variables. Individual MRIs were used for neuronavigation, but in one subject a separate more recent MRI was used in source modeling. The parameters of this specific MRI have not been included in the reporting. The Siemens MRI scanners were Skyra in Aalto and Prisma in Tübingen. 48 out of 50 had available T2-weighted MRIs. Some subjects also had diffusion tensor image and functional MRIs recorded, but these were not used in analysis.

| Parameter | Aalto / BioMag | Tübingen |
|---|---|---|
| **Participants** | | |
| Number of | 23[A] | 27 |
| Men/Women | 13/10 | 9/18 |
| Ages | 27.6 ± 6.9 | 27.5 ± 5.2 |
| rMT (% MSO) | 47.1 ± 7.8 | 51.7 ± 8.9 |
| **TMS** | | |
| Stimulator[B] | Nexstim NBT 2.2.4 / NBS 5.2.4 | Magventure R30/X100 |
| Coil (figure-of-eight) | Nexstim cooled | Cool-B65 |
| Neuronavigation | Nexstim NBT 2.2.4 / NBS 5.2.4 | Localite |
| ITI (s)[C] | 4.25 ± 0.25 | 3 ± 0.5 |
| Charge delay (s) | 1.9 | 1.0 |
| **EEG** (NeurOne) | | |
| Channels & layout | 64 & 10–20 | 128 & 10–5 |
| Reference location | Right mastoid | FCz |
| Ground location | Right cheekbone | AFz |
| Current | DC | DC |
| Filter (Causal; Hz) | LP 1250 | LP TESLA 1250 |
| Sampling frequency (Hz) | 5000 | 5000 |
| **MRI** (T1/T2) | | |
| Siemens scanner | Skyra | Prisma |
| Slice thickness (mm) | 1/1 | 0.8/0.8 |
| Repetition time (ms) | 2.53/3.2 | 2.4/3.2 |
| Echo time (ms) | 0.0033, 0.00337, or 0.00329/0.412 | 0.00222/0.563 or 0.564 |
| Flip angle (degrees) | 7/120 | 8/120 |

Four participants were excluded based on corrupt data, protocol deviations, or artifactual data and are not included in reporting. We note that the data of an individual subject was analyzed only once, however, a part of the measurement sessions were repeated for reasons of replicability or experimental irregularities. For all of these subjects except one, the repetition measurement session was included. We also note that the experiments have involved multiple personnel, which leaves the possibility for subjectivity in experimental performance. This affects, for example, the motor hotspot (and the corresponding coil position and orientation) and whether it was defined by also considering TEP resolutions or extracted movement of FDI or APB, which can lead to defined hotspots having E-field maximums outside of the primary motor cortex, or anatomical or MEP fidelity. However, the TMS-induced E-field is not a single point on the cortex and this work relies mostly on response variability rather than comparing mean response amplitudes across subjects. Thus, errors in stimulation locations or intensities (such as misestimating of the rMT, which can deviate due to the number of trials used, the muscle(s) observed, and MEP quality and legitimacy assessments), as long as sufficient response variability is present, do not contaminate our results. We also note that the rMT estimate of one subject was lowered by 1 due to estimate overshooting. Additionally, we note that the noise masking intensity was intermittently adjusted in the middle of the experiment if the subject notified that they could hear the TMS click.

### 2 TMS–EEG PREPROCESSING

Our preprocessing pipeline included several steps for handling contaminated channels and trials. We implemented an automated channel and trial rejection algorithm that assessed global and local noise levels, with separate adaptations for pre- and post-stimulus periods. Baseline corrections were applied using the pre-innervation window $[-100, -10]$ ms for EMG and $[-20-10]$ ms for post-stimulus EEG before SOUND and after pulse second pulse artifact interpolation. The post-stimulus data was was average referenced after removing bad channels, prior and after SOUND, SSP–SIR, and after the second baseline correction. The pre-stimulus data was average referenced after removing bad channels and after filtering. We also note that average referencing is re-applied prior to inverse modeling in the pre-stimulus data, which due to zero-mean removal, only affects the data due to small numerical imprecisions. We also performed re-analysis on the performance of the preprocessing (on a separate run, which can be affected by run-wise differences in algorithms) while focusing on of dropping of low-variability channels (after threshold adjustment) and trials.

We note that blocks, partial blocks, or trials were excluded from the data prior to the discussed preprocessing if smaller irregularities, such as noise masking problems, were noted. Blocks or partial blocks were not loaded into the structure while trials with noted noise masking problems in otherwise reliable blocks were excluded manually after loading the data. Trials with duplicate event identifiers were also dropped. A maximum number of 1200 trials was processed for each subject after this procedure as some subjects had additional trials recorded. We also note that for one subject, three trials without TMS were present in preprocessing, but they were automatically excluded based on our rejection criterion.

#### 2.1 Channel rejection

For channel rejection, we iteratively removed channels (until no more channels were rejected) whose standard deviation z-score exceeded ± 2.5 in a 2-Hz highpass filtered pre-stimulus period ($[-1045, -45)$ ms before stimulation) or ± 3 in the post-stimulus period ($[10, 150)$ ms after stimulation). When detecting bad channels from the pre-stimulus period, we applied adjusted thresholds for specific channel regions to account for ocular artifacts (later suppressed with ICA) or otherwise less activity (*e.g.* while very unlikely, substantially less high-amplitude ongoing oscillations, such as less occipital alpha leakage, could cause small deviation statistics if almost no ocular artifacts are present; re-analysis of the preprocessing script revealed that only 1 channel across all subjects was not rejected because of the lower threshold adjustment): increases of 1.5 for frontal pole and anterior frontal channels, and 0.75 for frontal channels (FF and F channel arrays). Additionally, we rejected channels based on their peak characteristics in the pre-stimulus period. For high-amplitude peaks (channels that have strong sudden sharp peaks), we used a prominence threshold of 70 $\mu$V and scale factor of 1, while for low-amplitude peaks (channels that are constantly very noisy), we used a prominence threshold of 2 $\mu$V and scale factor of 4. Channels were rejected if their peak counts exceeded mean + std*scale factor and if they showed an average of 3 or more high-prominence peaks across trials.

On average, 10.5 ± 6.5% of EEG channels were rejected and later reconstructed.

#### 2.2 Trial rejection

Trial rejection employed similar statistical criteria across the pre- and post-stimulus periods. Trials were rejected based on three main criteria: We excluded trials where the standard deviation z-score exceeded ± 1.5 within the pre-stimulus period ($[-1045, -45)$ ms, or exceeded ± 2.5 in the post-stimulus period $[10, 150)$ ms. We also rejected trials where the standard deviation z-score of at least one or two channels exceeded ± 6 or 5, respectively, in either time period. Finally, we removed trials with EMG activity where the peak-to-peak exceeded 50 $\mu$V in a baseline-corrected and linearly detrended window of 100–10 ms before stimulation. To optimize trial retention while maintaining data quality, we incrementally increased the EMG threshold by 10 $\mu$V (up to a maximum of 100 $\mu$V) until the EMG trial rejection rate fell below one-third of trials that were not rejected due to EEG. However, we note that this results in a higher effect of powerline noise on MEP amplitude estimates.

Based on these criterion, 14.4 ± 9.6% of trials were rejected on average. We note that absolute z-score thresholding on right-skewed distributions can mark valid trials with little variability as bad. Re-analysis of the preprocessing script revealed that on average, only 0.6 $\pm$ 2.3 trials were rejected because of having an overall low variability.

#### 2.3 Artifact suppression

We applied several artifact suppression techniques with parameters presented in Table 2.

**Table 2. Preprocessing parameters.** Independent component analysis (ICA; Hyvärinen and Oja (2000); Pion-Tonachini et al. (2019)) was fit to 2-Hz highpass filtered and average referenced pre-stimulus data in $[-1045, -45)$ ms compressed to 35 principal components and applied to average-referenced pre- and post-stimulus data to avoid data contamination and suppressing TMS-related neural activity. The source-estimate-utilizing noise-discarding (SOUND; Mutanen et al. (2018)) and the signal-space-projection-source-informed reconstruction (SSP–SIR; Mutanen et al. (2016)) algorithms were applied only to post-stimulus-focused EEG data. Filtering was only applied to pre-stimulus data. The post-stimulus EEG data was baseline corrected using data from $[-20, -10]$ prior to TMS. PCA: Principal component analysis.

| Method | Specification |
|---|---|
| ICA | Ocular artifact suppression using 35 PCA components where the IClabel probability of an ocular artifact exceeded 0.75 |
| SOUND | Regularization $(\lambda) = 0.05$ with 5 iterations |
| SSP–SIR | Removal of PCs explaining $\geq 90\%$ of variance in the supra 100-Hz data muscle artifact relative amplitude kernel estimated using 50-ms sliding windows |
| Filtering (Hz) | 2–90 bandpass and 48–52 bandstop, 4$^{\text{th}}$ order zero-phase Butterworth filters |
| Pulse artifact interpolation | Time windows of $[-5, 6]$ and $[-5, 8]$ ms relative to TMS with cubic interpolation using 1 ms of data before and after the respective time windows |

### 3 HEAD AND INVERSE MODELING

The cortical reconstructions were performed with FreeSurfer (Reuter et al., 2012) using T1-weighted MRIs with additional T2-weighting (T2 images were available for 48 out of 50 subjects) for pial surface contrast enhancement. The surface source spaces were subdivided into grade six octahedrons. We constructed three-layer head models with the boundary element method (BEM) with respective conductivities of 330, 4.2, and 330 mSm (Stenroos and Nummenmaa, 2016) between the boundaries of brain, inner and outer skull, and skin tissues that were reconstructed using the watershed algorithm (Ségonne et al., 2004), with additional corrections (homogeneously shrinking the inner skull surface) applied to resolve intersecting inner and outer skull surfaces. The forward solutions, *i.e.*, lead fields, describing source-sensor relationships (number of channels × 8196 sources * 3 orientations), were computed using these subject-specific source spaces, head models, and EEG–MRI co-registrations. Fiducials were also digitized, except for one subject, where they were retrieved from scalp landmarks. Faulty or non-existing individual electrode positions were identified by detecting outliers and visual inspection, and corrected using estimates based on coordinate transformations between the valid individual subject and the *standard_1005* electrode positions and manual adjustments.

The distributed source models were acquired with minimum-norm estimation using MNE Python (Gramfort et al., 2013; Hämäläinen and Ilmoniemi, 1994) with regularization $\lambda^2 = 0.1$, a depth weighting of 0.8, diagonal noise covariance with standard deviation of 0.2 $\mu$V, and data whitening with sources fixed normal to the cortical mantle.

##### 3.1 Average TEP characteristics

The resulting characteristics of dipoles fitted to the average response are displayed in Table 3.

**Table 3. Response characteristics across MEPs and dipolar TEP components fitted to the average response.** Values are presented as mean ± STD, except for window range. $N$: Number of included subjects whose dipole fitted to the average response had a coefficient of determination ($R^2$) of at least 75% and a dipole amplitude of at least 20 nAm. $R^2$ (%): Coefficient of determination of the dipole. Amplitude (nAm): Amplitude of the of the optimal dipole. Peak-to-peak ($\mu$V): Difference between minima and maxima of the sensor-space signal of the forward projected dipole or the MEP. Dipole latency (ms): Post-TMS time of the optimal dipole. Window size (ms): Width of the single-trial extraction window. Max window limits (ms): The min–max response fitting time across all subjects (constant for MEP). Note that the responses were determined in causal order individually for each subject. Source locations are given in MNI Talairach coordinates (mm). Unit-length orientations (from the native subject space) have been scaled by 100 for readability.

| **Parameter** | **N15** | **P30** | **N45** | **P60** | **MEP** |
|---|---|---|---|---|---|
| **Fit Overview**, $N$= | 11 | 23 | 41 | 45 | 50 |
| $R^2$ (%) | 84.4 ± 5.7 | 90.2 ± 6.1 | 88.9 ± 6.4 | 93.0 ± 5.0 | - |
| Amplitude (nAm) | 59.4 ± 41.7 | 48.6 ± 35.4 | 73.3 ± 84.4 | 61.3 ± 34.8 | - |
| Peak-to-peak ($\mu$V) | 9.0 ± 4.0 | 8.2 ± 5.1 | 10.9 ± 8.1 | 11.1 ± 6.3 | 1144 ± 1030 |
| Latency (ms) | 16.4 ± 1.7 | 33.0 ± 3.2 | 45.6 ± 3.0 | 63.5 ± 3.5 | - |
| Window size (ms) | 7.7 ± 1.4 | 5.1 ± 1.7 | 7.5 ± 2.3 | 12.2 ± 4.0 | - |
| Max window limits (ms) | 12.0–25.0 | 25.0–40.0 | 40.0–55.0 | 55.0–73.0 | 20.0–50.0 |
| **Location** | | | | | |
| x | 13.1 ± 19.2 | 5.0 ± 20.0 | -9.1 ± 17.0 | -6.7 ± 16.4 | - |
| y | -28.4 ± 26.0 | -22.6 ± 21.5 | -15.0 ± 27.7 | -20.3 ± 19.6 | - |
| z | 26.9 ± 26.8 | 38.7 ± 24.1 | 18.9 ± 26.0 | 31.2 ± 19.7 | - |
| **Orientation** | | | | | |
| x | 46.4 ± 32.7 | 2.0 ± 37.6 | -38.5 ± 38.1 | -43.9 ± 35.2 | - |
| y | 27.6 ± 34.8 | 11.6 ± 44.0 | -35.6 ± 33.5 | -42.4 ± 39.1 | - |
| z | 38.2 ± 57.9 | 55.7 ± 58.4 | -5.6 ± 68.2 | 0.5 ± 59.2 | - |

### 4 FEATURE EXTRACTION

#### 4.1 Feature extraction specifications

Due to measurement device offsets, jitters, and to avoid event mismatches, the neuronavigation coil control data was aligned with EEG by the smallest distances of real-world latencies after determining temporal offsets. Partially missing coil control data were replaced with the median of the existing deviations. In total, 4.7 ± 15.2% trials were replaced with the median of the individual subject. The across-subject average targeting accuracies of the preprocessed EEG trials were as follows, position: 2.2 ± 1.0 mm, direction: 1.3 ± 0.7, and normal: 1.3 ± 0.8 degrees. Additionally, in one subject, we used the median stimulation as the target after visual inspection of the original target location and stimulation points and one subject had completely missing coil control data, where the data was replaced with the median position, normal, and direction deviances across all subject means. In total, 6.2% of trials of the preprocessed EEG were handled with median replacement.

**Table 4. Feature extraction parameter specifications.** $N$ = number of time points, $f_s$ = sampling frequency. Phastimate parameters (excluding orders) are in samples (note that for this, the sampling rate $f_s = 1$ kHz). If no alpha peak was found, a default value of 10.5 Hz was used. The $f_{\alpha_{max}}$ values were 9.7 ± 1.2, range: 8.0–12.0 Hz, excluding 5 subjects where the default value was used. We positively shifted the coil control parameters by 1 and $10^{-3}$, respectively, to ensure successful log-transformations.

| Parameter | Specification |
|---|---|
| Bandwidth | $s_x f_s / N$, $s_\theta = 2, s_\alpha = 2, s_\beta = 3, s_\gamma = 6$ |
| PSD | multi-taper spectrum estimation |
| $\alpha$-phase | autoregressive model order = 25, edge = 65, Hilbert window length = 128, offset correction = 46 data window length = 1000, filter order = 192, and data cropped to maximum of 46 ms before TMS |
| $\alpha$-peak ($f_{\alpha_{max}}$) | detected from 7–14 Hz PSD with bandwidth $3f_s/2N$ using data $[-1045, -45)$ ms before TMS |
| Frequency range (Hz) | $\mathbf{f}_\alpha = f_{\alpha_{max}} \pm 2.5$, $\mathbf{f}_\theta = [\min(\mathbf{f}_\alpha) - 3.5, \min(\mathbf{f}_\alpha)]$, $\mathbf{f}_\beta = [\max(\mathbf{f}_\alpha), \max(\mathbf{f}_\alpha) + 20]$, and $\mathbf{f}_\gamma = [\max(\mathbf{f}_\beta), \max(\mathbf{f}_\beta) + 50]$ |
| Coil control | weighted mean + $10^{-3}$ of position (mm), normal (degrees) and direction (degrees) differences to the target, with respective weights = (0.70, 0.15, 0.15) |
| Time (trials) | Trial index of the preprocessed trials normalized by the number of maximum index of preprocessed trials |

### 5 PREDICTIVE MODELING AND STATISTICS

#### 5.1 Band-power–time and predictors

We found that the band-power–time-delay interaction parameters had significant but relatively smaller effect sizes compared to pure power-based predictors. The results for each anatomical parcel is displayed below.

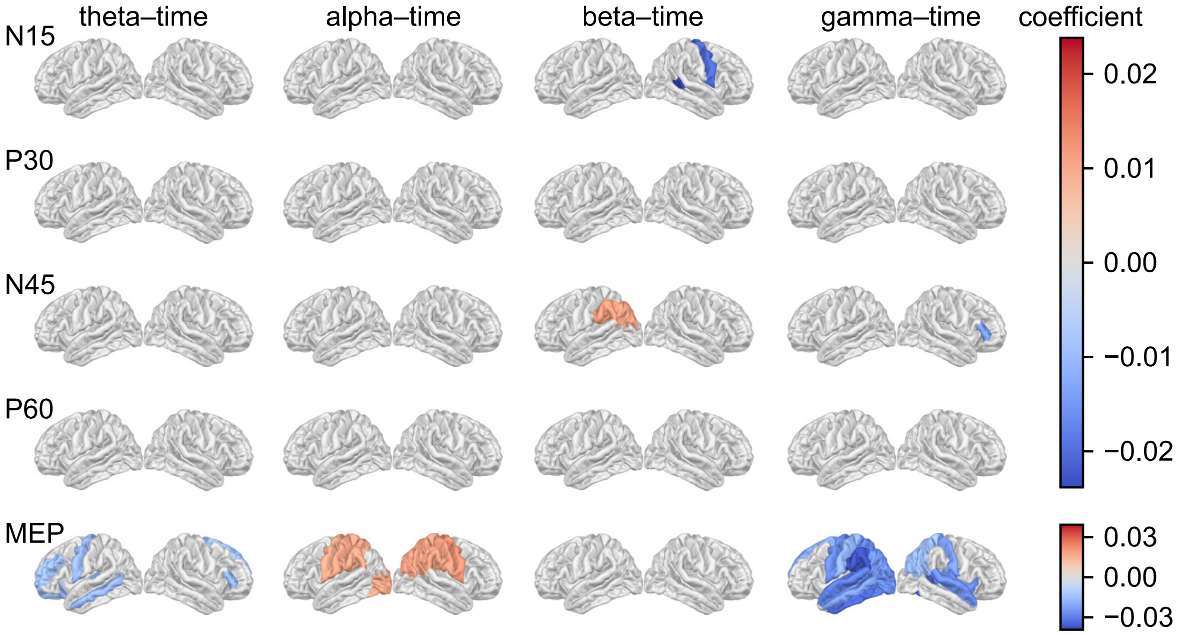


**Figure 1.** **Linear mixed-effect model analysis of band-power–time interaction.** Each anatomical parcel represents the frequency band-power coefficient of the model of the respective parcel modeling the respective response. Gray regions indicate insignificant ($p \geq 0.05$, Bonferroni corrected) coefficients. The coefficients are derived from the same models as in the main text.

We also fit models with alpha-phases estimated at each parcel at the time of TMS with *phastimate* (Zrenner et al., 2020). Here, trials where *phastimate* was not successful in prediction (resulting in NaNs) were dropped (separately for anatomical and custom parcels). We found that including phases in the models for anatomical parcels generally reduced model fidelity and it was thus not included in the final models. However, we acknowledge that we applied *phastimate* in a non-standard manner (no sensor-space Laplacian) in the source space and that we had multiple other predictors within the same model rather than only the phase decomposed to its sine and cosine components.

###### *5.1.1 Power–time interaction*

We found that band-power–time interaction effects had generally smaller but significant effects. This suggests that while there are temporal trends in frequency band activity, its effect on cortical and corticospinal responsiveness is generally smaller relative to instantaneous fluctuations in frequency band-power. The ongoing measurement time had a relatively larger effect on MEPs compared to TEPs, suggesting short-term modulations in muscular activity, such as fatigue, or corticospinal responsiveness.

#### 5.2 Marginal and conditional $R^2$

The marginal and conditional $R^2$, $R_c^2$, and $R_m^2$, respectively, were calculated for each model as

$$R_c^2 = \frac{\sigma_f^2 + \sigma_r^2}{\sigma_f^2 + \sigma_r^2 + \sigma_e^2} \tag{1}$$

and

$$R_m^2 = \frac{\sigma_f^2}{\sigma_f^2 + \sigma_r^2 + \sigma_e^2}, \tag{2}$$

where $\sigma_f^2$, $\sigma_r^2$, and $\sigma_e^2$ are the variances of fixed effects, random effects, and residuals, respectively (Nakagawa and Schielzeth, 2013).
